# Probabilities of Fitness Consequences for Point Mutations Across the Human Genome

**DOI:** 10.1101/006825

**Authors:** Brad Gulko, Melissa J. Hubisz, Ilan Gronau, Adam Siepel

## Abstract

We describe a novel computational method for estimating the probability that a point mutation at each position in a genome will influence fitness. These *fitness consequence* (fitCons) scores serve as evolution-based measures of potential genomic function. Our approach is to cluster genomic positions into groups exhibiting distinct “fingerprints” based on high-throughput functional genomic data, then to estimate a probability of fitness consequences for each group from associated patterns of genetic polymorphism and divergence. We have generated fitCons scores for three human cell types based on public data from ENCODE. Compared with conventional conservation scores, fitCons scores show considerably improved prediction power for *cis*-regulatory elements. In addition, fitCons scores indicate that 4.2–7.5% of nucleotides in the human genome have influenced fitness since the human-chimpanzee divergence, and, in contrast to several recent studies, they suggest that recent evolutionary turnover has had limited impact on the functional content of the genome.

---

During the past decade, two major developments—the emergence of massively parallel, ultra-cheap DNA sequencing technologies and the use of these technologies as digital readouts for functional genomic assays—have led to a profusion of data describing various features of genomes, epigenomes, and transcriptomes^1, 2^. However, investigators still have only rudimentary tools for integrating these diverse sources of information to obtain useful insights about genomic function and evolution. The limitations of current methods are particularly evident in the vast noncoding regions of eukaryotic genomes, which, despite important recent progress^3–6^, remain poorly annotated and understood. These limitations hamper progress in many areas, ranging from basic molecular genetics to disease association and personalized medicine^7^.

Many computational methods for gaining functional insights from sequence data are based on the simple, but powerful, observation that functionally important nucleotides tend to remain unchanged over evolutionary time, because mutations at these sites generally reduce fitness and are therefore eliminated by natural selection^7–15^. A major strength of these conservation- or constraint-based approaches is that they sidestep thorny questions about the relationship between the outcomes of biochemical experiments and fitness-influencing functional roles^16–19^ by getting at fitness directly through observations of evolutionary change. In essence, the “experiment” considered by these methods is the one conducted directly on genomes by nature over millenia, and the outcomes of interest are the presence or absence of fixed mutations. These conservation-based methods, however, depend critically on the assumption that genomic elements are present at orthologous locations and maintain similar functional roles over relatively long evolutionary time periods. Evolutionary turnover may cause inconsistencies between sequence orthology and functional homology that substantially limit this type of analysis.

This important limitation has led to two major alternative strategies for the identification and characterization of functional elements. The first strategy is to augment information about interspecies conservation with information about genetic polymorphism^20–28^. The shorter evolutionary time scales associated with intraspecies variation make this approach more robust to evolutionary turnover and less sensitive to errors in alignment and orthology detection. Polymorphic sites tend to be sparse along the genome, however, so this approach requires some type of pooling of information across genomic positions, which can be difficult in the absence good-quality genomic annotations. The second strategy is to forgo the use of evolutionary information and instead to predict functional roles from genomic data alone, typically with machine-learning methods for supervised classification^29, 30^ or clustering followed by labeling based on known examples^31–33^. This approach has the limitation that it depends strongly on previously characterized elements, which, in noncoding regions, are typically small in number and perhaps unrepresentative of the genome.

In this paper, we introduce a method for genomic analysis that combines many of the strengths of these polymorphism-based and functional genomic approaches. Like functional genomic methods, our approach groups genomic regions according to functional genomic “fingerprints” across multiple assays. Instead of relying on known examples for classification, however, we characterize each group by a probability of mutational fitness consequences—or fitCons score—inferred from patterns of genetic variation. These fitCons scores are estimated using a recently developed statistical method, called Inference of Natural Selection from Interspersed Genomically coHerent elemenTs (INSIGHT), that contrasts patterns of polymorphism and divergence in a collection of dispersed genomic sites with those in nearby neutral sites, accounting for negative and positive selection^34^. Thus, the method integrates both evolutionary and functional data in characterizing the potential functional importance of genomic regions. We demonstrate that these fitCons scores are useful for visualization, for prediction of *cis*-regulatory elements, and for measuring the global influence of recent natural selection across the genome.

## Results

### General Features of the Prediction Problem

Information about genetic variation can be used to estimate probabilities of fitness consequences for moderately large groups of genomic positions but not for individual loci, owing to the sparsity of informative sites along the genome. This property of “groupwise” but not “individual” predictivity is common to many statistical problems, but it is complicated in our case by two additional features. First, an appropriate scheme for grouping or stratification is not clear *a priori* here. Second, the outcome of interest in our problem— mutational fitness consequences—is not directly observable from the data. For contrast, consider the problem of estimating the expected risk of an automobile accident. This problem must also be addressed at the level of groups (either explicitly through stratification of drivers, or implicitly through regression), but in this case, the relevant features—such as the age, sex, and number of traffic violations of the driver—are obvious. In addition, the outcomes of interest—the occurrence and cost of accidents—are directly observed. In our problem, the genomic “risk factors” for fitness-influencing mutations, particularly in unannotated noncoding regions of the genome, are much less clear. Furthermore, once a grouping is determined, it is still not possible to read off the associated fitness consequences of mutations; instead they must be inferred from patterns of genetic variation using an evolutionary model.

### Calculation of FitCons Scores

We have addressed these challenges using the following strategy. Beginning with genome-wide functional genomic data sets obtained from each cell type (Fig. 1A), we first cluster genomic positions by their joint functional genomic “fingerprints” (Fig. 1B). We expect three highly informative functional genomic data types—DNase-seq, RNA-seq, and ChIP-seq data describing histone modifications—to represent largely orthogonal sources of information, describing DNA accessibility, transcription, and chromatin states, respectively. Therefore, we divide genomic positions into three levels of DNase-seq “signal,” depending on whether they fall outside of ENCODE-designated DNase-seq peaks (0), in broad peaks only (1), or in narrow peaks (2); into four groups having no aligned RNA-seq reads (0) or low (1), medium (2) or high (3) mean RNA-seq read depth; and into 26 distinct chromatin states categories based on the ChromHMM method^31, 33^. In addition, we distinguish between sites that fall outside (0) or within (1) annotated protein-coding sequences (CDSs), which we expect to show pronounced differences in selective pressure. We then consider all possible combinations of these four types of assignments, obtaining 3 *×* 4 *×* 26 *×* 2 = 624 distinct functional genomic classes. This clustering step was applied separately to three karyotypically normal cell types: human umbilical vein epithelial cells (HUVEC), H1 human embryonic stem cells (H1 hESC), and lymphoblastoid cells (GM12878), resulting in 443–447 usable classes of sites, with median numbers of 165 to 224 thousand sites per class (see Supplementary Table S1 and Methods for details).

**Figure 1:**
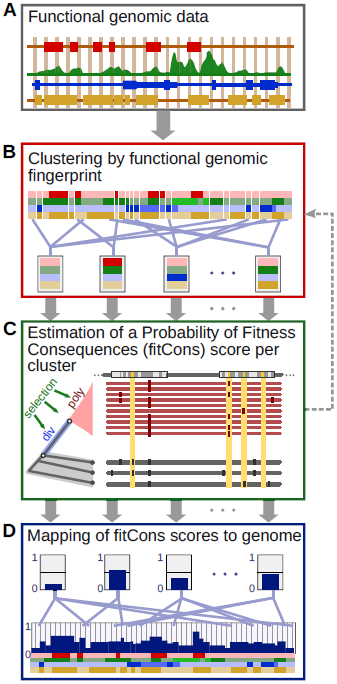
Illustration of procedure for calculating fitCons scores. (A) Functional genomic data, such as DNase-seq, RNA-seq and histone modification data, are arranged along the genome sequence in tracks. (B) Nucleotide positions in the genome are clustered by joint patterns across these functional genomic tracks. For example, one cluster might contain genomic positions with a high DNase-seq signal, a moderate RNA-seq signal, and high signals for H3K4me1 and H3K27ac, suggesting transcribed enhancers. Another might contain positions with a low DNase-seq signal, a high RNA-seq signal, and a signal for H3K36me3, suggesting actively transcribed gene bodies. Notice that clusters will generally contain genomic positions dispersed along the genome sequence. (C) Patterns of polymorphism and divergence are analyzed using INSIGHT^34^ to obtain an estimate of the fraction of nucleotides under natural selection (*ρ*) in each cluster. This quantity is interpreted as a probability that each nucleotide position influences the fitness of the organism that carries it, or a fitness consequence (fitCons) score. (D) The fitCons score for each cluster is assigned to all genomic positions that were included in the cluster. In this way, all nucleotide positions are assigned a score, but there can be no more distinct scores than there are clusters. Note that, in our initial work, the clustering is of genomic positions is accomplished by a simple exhaustive partitioning scheme that produces 624 distinct clusters. In future work, however, it may be desirable to iterate between clustering and calculating scores (dashed line; see Discussion).

Next, we use INSIGHT to estimate the probabilities of mutational fitness consequences within each of these classes based on patterns of polymorphism and divergence (Fig. 1C). This step yields an estimate of *ρ* for each of the analyzed classes, which serves as the fitCons score for that class. Finally, we assign to each nucleotide position in the genome the score estimated for the corresponding functional genomic class (Fig. 1D). Each genomic position is thus assigned a value between 0 and 1, representing the probability that the nucleotide at that position influences fitness, as estimated from patterns of variation at all genomic sites displaying the same functional genomic fingerprint. These fitCons scores uniquely integrate information from evolutionary and cell-type-specific functional genomic data.

### Genomic Distribution of FitCons Scores

To obtain a general overview of the genomic distribution of fitCons scores, we first considered the *composition* and *coverage* of nucleotide sites of various annotation types as the fitCons score threshold was varied, focusing on HUVEC (see Discussion for other cell types). When the score threshold *S* = 0, all sites are included and the composition of annotations reflects the overall genomic distribution (Fig. 2A). As *S* increases, however, the sites in known functional classes become strongly enriched relative to the intergenic and intronic sites. Regions such as 5′ and 3′ UTRs, promoters, and introns are most enriched at intermediate scores, reflecting moderate levels of natural selection in these regions, while CDSs dominate at the highest scores. The coverage properties (Fig. 2B) are best for CDSs, 3′ UTRs, and 5′ UTRs (in that order), but they are also considerably elevated above the intergenic background for promoters, transcription factor binding sites (TFBS), long noncoding RNAs (lincRNAs), and small noncoding RNAs (snRNAs). Notably, the enrichment for functionally annotated genomic regions at high scores occurs despite no use of genomic annotations in the scoring scheme (with the single exception of CDS annotations). Instead, these elevated scores reflect differences in patterns of polymorphism and divergence that arise naturally from the fitness consequences of mutations in these regions.

**Figure 2:**
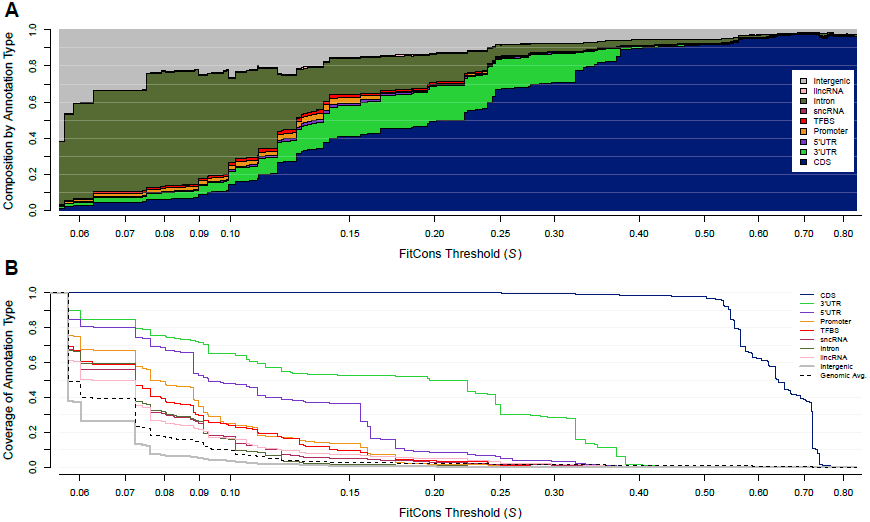
Composition and coverage of high-scoring genomic regions according to fitCons. (A) Composition by annotation type in regions that exceed a fitCons score threshold of *S*, as *S* is varied across the range of possible scores. Each vertical cross-section of the plot can be thought of as a narrow “stacked bar” representation of the composition by annotation type of all genomic positions at which the fitCons score > *S*. At the left side of the plot, when *S* is small, the composition by annotation type is representative of the genome as a whole. As the threshold *S* increases, coding sequences (CDS; navy) are increasingly enriched and intergenic (gray) sequences are increasingly depleted. Regions experiencing moderate levels of selection, such as untranslated regions (UTRs; purple and green), promoters (orange), small noncoding RNAs (sncRNAs; eggplant), and introns (olive), are most enriched at intermediate scores. A logarithmic scale is used for the *x*-axis to enable the differences at low score thresholds to be observed more clearly. (B) Coverage of the same annotation types by genomic regions having fitCons score > *S*, with an *x*-axis matching that in (A). The dashed line indicates the genome-wide average. At each value of *S*, the relative height of a given curve in comparison to the dashed line indicates the enrichment (or depletion) of the corresponding annotation type in genomic regions having score > *S*. The legend at the right lists the annotation types in order of decreasing enrichment.

We have developed UCSC Genome Browser tracks that display fitCons scores across the genome for each cell type of interest. FitCons scores are clearly elevated in CDS and UTR exons, like existing evolutionary conservation scores, but they are often better at highlighting candidate regulatory elements, particularly outside the core promoter (Fig. 3). Many of these regulatory elements may not be conserved across species because they have arisen recently in evolutionary time, while others may have spuriously low conservation scores because of errors in orthology detection or alignment. In agreement with anecdotal observations from the Genome Browser, the fitCons scores are fairly well correlated with phyloP conservation scores^15^ genome-wide, with some notable exceptions in noncoding regions (Supplementary Fig. S1).

**Figure 3:**
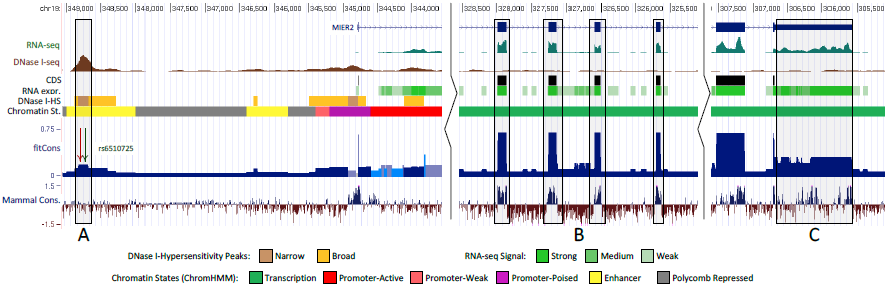
Genome browser display showing functional genomic fingerprints and fitCons scores. Shown, from top to bottom, are the exons of the *MIER2* gene (blue); the raw RNA-seq (turquoise) and DNase-seq (brown) signals; the four discretized tracks used to define the 624 functional genomic fingerprints, including annotation-based CDSs (black), RNA-seq signal (green), DNase-seq signal (yellow and brown), and chromatin modifications (multiple colors; see key at bottom); the fitCons scores based on those fingerprints (dark blue, with lighter blues less statistically significant); and, for comparison, phyloP-based conservation scores for mammals. (A) An apparent enhancer, marked by a combination of enhancer-associated chromatin modifications and a strong DNase-seq signal, displays elevated fitCons scores but no elevation in conservation scores. A ChIP-seq-supported TFBS for AP-1 (red arrow) and a lung-cancer-associated SNP (green arrow) are highlighted. (B) CDS exons show elevated scores according to both fitCons and phyloP. (C) The 3′ UTR, marked by transcription-associated chromatin modifications, a high RNA-seq signal, and an absense of DNase-I hypersensitivity or CDS annotations, displays moderately elevated fit-Cons scores and patches of evolutionary conservation. Browser tracks are publicly available at http://genome-mirror.bscb.cornell.edu (hg19 assembly).

The fitCons scores tend to increase with the strength of the DNase-seq signal (Fig. 4A), probably due to an increasing density of *cis*-regulatory elements. They also increase with the RNA-seq signal (Fig. 4B), evidently because of changes in the type of transcription unit identified (e.g., non-CDS regions with high RNA-seq signal are enriched for relatively conserved 3′ UTRs). When joint patterns of DNase-seq and RNA-seq are considered, however, a more complex pattern emerges: fitCons scores increase with DNase-seq intensity at low RNA-seq intensities, as they do overall, but this trend is reversed at high RNA-seq intensities (Fig. 4C). A closer examination reveals that this pattern is explained by the enrichment for DNase-I-hypersensitivity near the 5′ ends of genes. In particular, conditional on a high RNA-seq signal, a high DNase-seq signal tends to be associated with 5′ UTRs and upstream regions of genes, which are under somewhat weaker selection than the 3′ UTRs associated with lower DNase-seq signals. This example demonstrates that the fitCons scores can capture non-additive and potentially surprising relationships between functional genomic covariates and natural selection.

**Figure 4:**
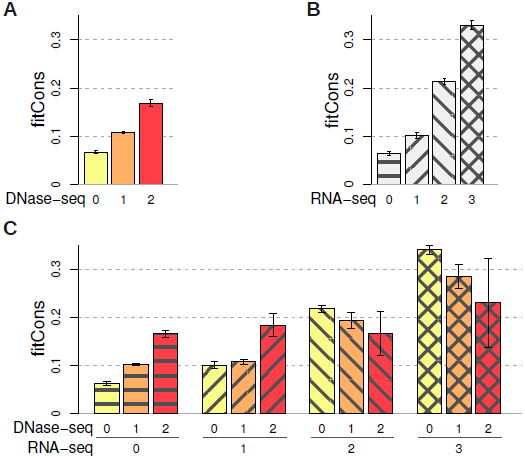
Average fitCons scores as a function of DNase-seq and RNA-seq intensity. Results represent averages across all non-CDS clusters having the marginal or joint property of interest. Error bars represent standard errors of the aggregated fitCons scores (see Methods). (A) FitCons scores increase with DNase-seq intensity. 0: No DNase-seq signal; 1: broad peaks; 2: narrow peaks. (B) FitCons scores increase with RNA-seq intensity. 0: no RNA-seq reads; 1-3: weak to strong RNA-seq signal (see Methods). (C) FitCons scores behave in a non-additive manner as joint combinations of DNase- and RNA-seq intensity are considered. In particular, at medium to high RNA-seq read depth (classes 2 and 3), fitCons scores decrease (rather than increase) with increasing DNase-seq signal.

### Predictive Power for *Cis*-Regulatory Loci

We evaluated the predictive power of the fitCons scores for known cell-type-specific regulatory elements in comparison with three widely used phylogenetic conservation scoring methods, the phastCons^12^, phyloP^15^ and Genomic Evolutionary Rate Profiling (GERP)^13^ programs. In addition, we considered a new program, called Combined Annotation Dependent Depletion (CADD)^35^, that estimates relative levels of pathogenicity of potential human variants using a support vector machine (SVM), many different genomic annotations, and simulations of nucleotide divergence rates. Where possible, we also considered RegulomeDB, a scoring system for the regulatory potential of variant sites based on combined experimental and computational data^36^. We evaluated the performance of these scoring methods in predicting three types of functional elements with putative roles in transcriptional regulation: binding sites for various transcription factors supported by chromatin immunoprecipitation and sequencing (ChIP-seq) data from the ENCODE project^3, 28^; (2) high-resolution expression quantitative trait loci (eQTL) identified in a recent large-scale study^6^; and (3) enhancers based on characteristic chromatin marks^37^ (see Methods for details). Importantly, these annotation sets are based on different functional genomic data from that used by fitCons, with the exception of some overlap between the data used in set (3) and that used in our classification of chromatin states.

To place the different scoring methods on equal footing, we plotted the basewise coverage of each type of regulatory element as a function of the total coverage of the noncoding genome, varying score thesholds to include 0–20% of noncoding sites (Fig. 5). This strategy allows us to measure the extent to which the elements of interest display “signals” that rise above the “noise” of the noncoding genome, in a uniform manner across scoring methods. By this test, the fitCons scores showed dramatically better sensitivity for noncoding elements than all of the other methods considered. For example, at a total noncoding coverage of 2.5%, fitCons scores achieve nearly 70% coverage of TFBSs, whereas the other methods all have less than 15% coverage. Similarly, the coverage of enhancers was about 40% at 2.5% noncoding coverage, while the other scoring methods showed almost no signal above background. We also performed a more traditional evaluation of the trade-off between sensitivity and specificity using receiver operating characteristic (ROC) curves and found that, in all cases, fitCons scores were considerably better predictors of regulatory function than phastCons, phyloP, GERP, CADD, or RegulomeDB scores (see Methods and Supplementary Fig. S2).

**Figure 5:**
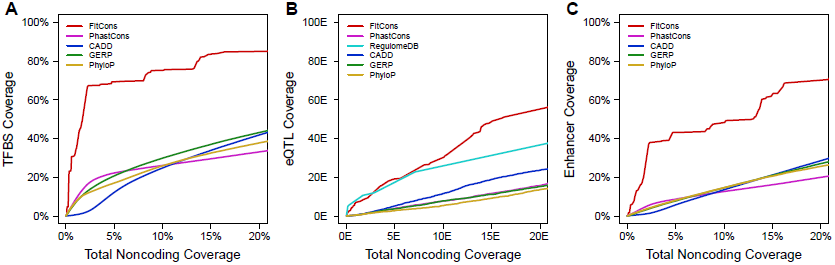
Coverage of active *cis*-regulatory elements as a function of total coverage of the noncoding genome. Coverage of each type of element is shown as the score threshold is adjusted to alter the total coverage of noncoding sequences in the genome, excluding sites annotated as CDS or UTR. FitCons is compared with scores from the Combined Annotation Dependent Depletion (CADD)35, Genomic Evolutionary Rate Profiling (GERP)13, phastCons12, and phyloP15 programs (see Methods). (A) Coverage of transcription factor binding sites detected by ChIP-seq in HUVEC28. (B) Coverage of high-resolution eQTLs identified in a recent large-scale study6, restricted to eQTLs associated with genes transcribed in HUVEC. Coverage of eQTLs is also shown for classification of single nucleotide variants by RegulomeDB36. The divergence-based scores (phastCons, phyloP, GERP, and CADD) all perform poorly on the eQTL data set, probably because the ascertainment for segregating sites creates a bias against evolutionary conservation. Note also that the apparent performance of RegulomeDB, particularly at low total noncoding coverage, is somewhat influenced by consideration of eQTL data in its scoring scheme. (C) Coverage of enhancers identified by characteristic chromatin marks37 assayed in HUVEC.

The tests above were based on regulatory elements that are putatively active in the cell type for which the scores were produced, to exploit fitCons’s use of cell-type-specific functional data. To evaluate how well these advantages carried across cell types, we created an “integrated” fitCons score by combining information from three cell types (see Methods), and evaluated the performance of this score in predicting regulatory elements pooled from multiple cell types. We found that, in this less favorable setting, the fitCons scores still had better predictive performance for *cis*-regulatory elements than any of the other scoring methods (Supplementary Fig. S3). In addition, we carried out a second set of tests of ChIP-seq-supported TFBSs that considered only the subset of nucleotide positions at which base preferences were especially strong, which should be enriched for sites at which mutations will have fitness consequences. The ROC curves based on this more stringent test were very similar to the original curves (Supplementary Fig. S4), demonstrating that the apparent performance of the fitCons scores was not artificially inflated by the inclusion of neutral sites in our validation experiments and the relatively coarse resolution of the scores.

### Proportion of the Human Genome Under Selection

The proportion of nucleotides in the human genome that directly influence fitness—sometimes called the “share under selection” (SUS)—has primarily been estimated using methods that consider divergence patterns among mammals, for which turnover of functional elements may be an important confounding factor^38–42^. The fitCons scores—in addition to being useful as predictors of function—could be useful in obtaining estimates of the SUS that are less sensitive to turnover because they measure natural selection over much shorter time scales.

An initial estimate of the SUS can be obtained by simply averaging the fitCons scores across all nucleotide positions in the genome. Because each score represents a probability that an individual nucleotide influences fitness, their average represents an expected fraction of nucleotides in the genome with fitness-influencing functions, or an expected SUS. This approach yields an estimate of 7.5% (*±*0.1%) for HUVEC, or 7.5–7.8% across cell types. Among sites under selection, we estimate that 9.0% are in CDS, 2.2% in 3′ UTRs, 35.2% in introns, 51.7% in intergenic regions, and *<*1% in each of several other noncoding annotation classes (Supplementary Table S2). These estimates are generally consistent with, but on the high end, of those based on cross-species divergence, which generally have fallen between 3 and 8% and have suggested that noncoding bases outnumber coding bases by factors of 2.5–8^12, 38, 42–44^. Interestingly, our estimates are somewhat lower than estimates that have explicitly allowed for evolutionary turnover, which have been 2–3 times higher than the pan-mammalian estimates of *∼*5%^26, 42, 44–46^ (see Discussion).

Furthermore, violations of modeling assumptions will tend to bias the fitCons scores upward, particularly for functional classes for which the true fraction is close to zero (see Supplementary Note). To address this problem, we performed a parallel calculation for “neutral” sites that intersect the large class of genomic positions having a “null” functional genomic fingerprint (i.e., no DNase-seq, RNA-seq, or histone modification signal). This results in an estimate of 3.3%, which can be considered an upper bound on the contribution of error because these putatively neutral sites undoubtedly include some sites under selection. By subtracting this 3.3% from our naive estimate of 7.5%, we obtain an estimated lower bound for the SUS of 4.2%, with somewhat higher fractions of selected sites in CDSs and 3′ UTRs (Supplementary Table S2). (These estimates are for HUVEC, but the results for the other cell types were very similar.) Overall, our analysis of the SUS suggests that between 4.2% and 7.5% of nucleotides in the genome have direct fitness-influencing functions, and that the ratio of noncoding to coding functional sites is between 5.4 and 10.1, estimates that are remarkably consistent with those based on measures of divergence between mammalian genomes (see Discussion).

### Implications for Evolutionary Turnover of Functional Elements

To better understand the differences between our new scores and scores based on divergence, we devised a set of alternative scores (denoted “fitConsD”) based on the same clusters of sites but an estimator of the fraction of nucleotides under selection that instead considers nucleotide divergence across primates (see Methods). Thus, the fitCons and fitConsD scores both represent probabilities of fitness consequences per nucleotide but over two different evolutionary time scales. Overall, these two measures are remarkably well correlated, with *R*^2^ = 0.88 (Fig. 6A). Under both scoring systems, the scores for coding sequences are generally higher than those for noncoding sequences, as expected. However, the noncoding sites exhibit greater variance (*R*^2^ = 0.51 vs. *R*^2^ = 0.69 for coding), and a slight excess of clusters with higher fitCons than fitConsD scores. These observations suggest that the main signal for selection has been maintained over long evolutionary time periods, but that there are some classes that show stronger recent than ancient natural selection.

**Figure 6:**
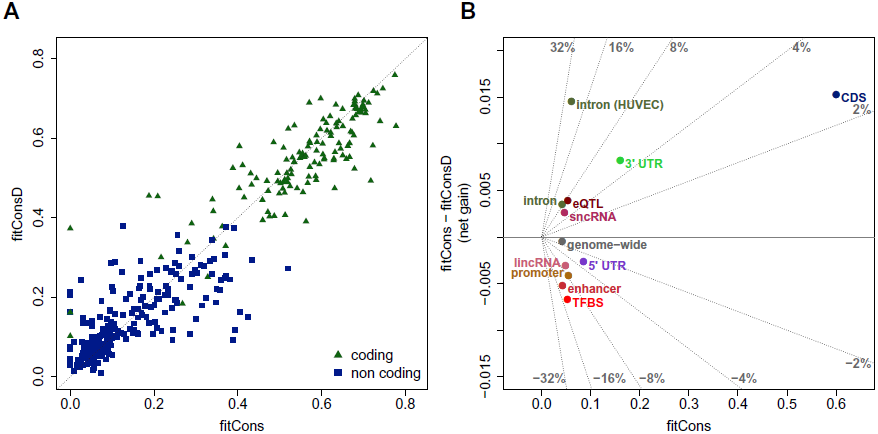
Comparison between fitCons and fitConsD scores. FitConsD is an alternative estimate of fitness consequences, analogous to fitCons, but based on an estimator of the fraction of sites under natural selection that considers divergence patterns across four primate genomes (see Methods). (A) FitCons and FitConsD scores are shown for the clusters defined using functional genomic data from HUVEC. Scores are shown for the 348 clusters of size 10 kb or larger, distinguishing between coding clusters (green triangles) and non-coding clusters (blue squares). Both sets of scores are corrected by subtracting the possible contribution from model misspecification (see Methods). Correlation between the two sets of scores is high (*R*^2^ = 0.88) overall, and is somewhat higher for coding (*R*^2^ = 0.69) than for noncoding (*R*^2^ = 0.51) clusters. (B) The net gain in the fraction of sites under selection on population genetic time scales relative to primate-divergence time scales, computed by subtracting average fitConsD scores from average fitCons score for different classes of functional elements. Net gain is plotted against average fitCons score, and lines of constant slope radiating from the origin represent constant values of a “net gain rate,” computed as NGR = (fitCons – fitConsD)/fitCons. The NGR can be interpreted as an estimate of the net gain per functional site. Notice that the NGR is small (≤10%) for almost all annotation classes considered. The main exception is driven by a few clusters associated with an absence of DNase-seq or RNA-seq signal and chromatin modifications suggesting transcriptional elongation, which had net gain rates as high as 20% and were strongly enriched in the introns of active genes (see “intron (HUVEC)”), suggesting the possibility of evolutionary innovation associated with mRNA splicing or post-transcriptional regulation.

The impact of turnover can be assessed more directly by subtracting the average fitConsD score from the fitCons score for various annotation classes of interest. This difference between average scores can be interpreted as the “net gain” in functional nucleotides on population genetic time scales relative to primate-divergence time scales, with a negative value representing a net loss (see Fig. 6B). This net gain can be further divided by the fitCons score for each annotation class of interest to obtain a net gain rate per functional nucleotide (diagonal lines in Fig. 6B). We observe relatively low amounts of turnover overall, with net gains for our standard set of annotation classes ranging between −0.7% and 1.7% and net gain rates roughly between −10% and 10%. These observations suggest that the net gain or loss of functional nucleotides in both coding and non-coding functional elements accounts for no more than about 10% of all functional sites, although these estimates do not exclude the possibility that offsetting gains and losses produce considerable evolutionary flux in the functional composition of the genome (see Discussion).

## Discussion

We have presented a new approach that combines functional genomic data with data describing variation within and between species. The essential idea of the approach is to use the functional genomic data to group sites into classes that are relatively homogeneous in terms of their functional roles, then to characterize the bulk influence of natural selection on those classes based on their patterns of polymorphism and divergence. For our estimation of natural selection, we make use of a recently developed probabilistic model of evolution and efficient algorithms for genome-wide inference (INSIGHT). We interpret INSIGHT-based estimates of fractions of nucleotides under selection as probabilities that each nucleotide influences fitness, or *fitness consequence* (fitCons) scores. Even with a simple clustering scheme, these fitCons scores appear to be highly informative about genomic function.

Based on our experiments, the fitCons scores have excellent predictive performance for putative *cis*-regulatory elements, outperforming several divergence-based methods (phastCons, phyloP, GERP, and CADD) and one annotation-based method (RegulomeDB) by clear margins. In part, this improvement in performance reflects the use of cell-type-specific data and, indeed, it is most pronounced when considering elements active in the cell-type used to compute the scores (Fig. 5 and Supplementary Fig. S2). Nevertheless, the fitCons scores also show a clear performance advantage when considering all annotated elements rather than just active ones (Supplementary Fig. S3). The approach of grouping genomic sites by functional genomic signatures, and then measuring groupwise fitness consequences based on patterns of genetic variation, appears to offer real benefits for prediction of regulatory function, as compared with methods that consider either genetic divergence or functional genomic data alone. In addition, in at least some cases, the use of population genomic data helps to identify elements that appear to have emerged recently in evolutionary time and therefore display no significant reductions in interspecies divergence.

Interestingly, the recently published CADD method performed no better on our tests than conventional conservation scores, despite reports by the authors of significant advantages over phyloP, phastCons, GERP, and other methods^35^. This difference in CADD’s performance appears to reflect several important distinctions between our validation experiments and those reported in reference [35]. First, our tests focused specifically on putative *cis*-regulatory elements, while many of their tests considered a mixture of coding and noncoding elements, in some cases, implicitly enriched for coding regions. In particular, the ClinVar database, which figured prominently in their experiments, includes very few noncoding variants (*∼*5% of pathogenic variants). Our results suggest that CADD may perform considerably better in coding regions than in *cis*-regulatory regions—not surprisingly, given the method’s use of numerous features derived from annotations of protein-coding genes. Second, when Kircher et al. did consider noncoding regions, they generally did not distinguish between *cis*-regulatory mutations and mutations that more directly influence the structure and content of protein-coding transcripts, such as mutations to splice sites or UTRs. CADD has a natural advantage with these variants also due to its consideration of gene annotations, whereas the annotation-free fitCons scores may perform better in completely unannotated regions of the genome. Finally, the tests by Kircher et al. that explicitly considered putative *cis*-regulatory elements were limited to a few loci and considered only correlations with saturation mutagenesis experiments, irrespective of a prediction threshold. We view our ROC-type comparisons based on three separate genome-wide sets of elements as a more direct and comprehensive demonstration of predictive power in *cis*-regulatory elements. It is worth noting that the goal of CADD is stated slightly differently from ours—its authors aim to predict pathogenicity of alleles rather than probabilities of fitness consequences at nucleotide positions—but our measure of fitness consequences and their measure of pathogenicity are essentially the same (evidence of negative selection from genetic variation), so we believe it is appropriate to compare the two methods directly. In any case, the comparison of these two closely related, yet distinct, approaches helps to reveal strengths and weaknesses of each of them, and may lead to new ideas for improved methodologies.

A side benefit of our model-based approach is that the basewise probabilities of fitness consequences lead in a straightforward manner to an estimate of the “share under selection” (SUS) in the human genome. This estimate of the SUS reflects time scales since the divergence of humans and chimpanzees, about 4–6 million years ago, unlike conventional estimates based on tens or hundreds of millions of years of mammalian evolution. Nevertheless, our estimate of the SUS—at 4.2–7.5%—ends up being remarkably similar to those based on longer time scales, which have generally fallen between 3 and 8%^12, 38–41, 43, 47^. We take the general concordance of these estimates, both with one another and with our fitCons- and fitConsD-based estimates, as an indication that the SUS has been fairly stable at roughly 5% over various time-scales in mammalian evolution. This finding stands in contrast to estimates of much higher functional contents in the genome, of 80% or more, based on measures of “biochemical activity”^3^. However, it is important to bear in mind that these evolutionary and biochemical estimates reflect somewhat different definitions of “function,” and this may explain some of the difference between them^16, 18, 19^. For example, the fitCons- and conservation-based estimates of the functional content of the genome generally represent fractions of positions at which a point mutation that alters the identity of a nucleotide will have fitness consequences, but they do not account for sequences (such as spacer elements) that would have fitness consequences if deleted but not mutated (see Supplementary Note for discussion).

Apart from the absolute fraction of functional DNA in the human genome is the question of how much the functional content of the genome has changed over time through gains and losses of functional elements. Several studies have attempted to estimate the impact of such “turnover,” and have concluded that the current SUS in the human genome could be *∼*2–3 times larger than estimates based on comparisons across mammals^26, 42, 44–46^. Indeed, these findings have been proposed to explain, in part, the discordance between evolution-based and biochemical estimates of the functional fraction of the genome^26, 48, 49^. However, these analyses have accounted for turnover using relatively crude methods, for example, by relying on an apparently near-linear relationship between pairwise divergence and the estimated SUS^44, 45^ or by estimating functional content from mean SNP densities or derived allele frequencies in genomic regions not conserved across mammals^26^. Our analysis is more direct, by comparing analogous divergence-based and polymophism-based estimates of the SUS based on exactly the same clusters of nucleotide positions. In addition, our analysis focuses on the more restricted question of evidence for selection on the time scales of primate evolution, rather than attempting to account for turnover across the mammalian phylogeny, where factors such as alignment error, orthology detection, and genomic rearrangements can be problematic. The similarity between our estimates based on polymorphism (fitCons) and divergence (fitConsD) strongly suggests that evolutionary turnover has been modest during primate evolution, because massive turnover would be expected to lead to a substantial downward bias in the divergence-based estimates. Furthermore, our power experiments suggest that this observation is not an artifact of reduced sensitivity in the fitCons scores. Nevertheless, we cannot rule out the possibility that compensating gains and losses on very recent time scales maintain a similar SUS while substantially altering the genomic composition of functional sequences.

An obvious area for improvement in our current methods is our approach to clustering. Our strategy of considering the Cartesian product of discrete covariate categories will not scale to large numbers of covariates. In addition, in some cases it results in overly coarse-grained clusters, whereas in others it fails to group together small clusters with similar fitness consequences. An iterative approach to clustering (Fig. 1) could enable the estimated fitCons scores to help guide the clustering scheme. This approach would be computationally intensive because of the expense of estimating model parameters from genome-wide data and the need for some type of regularization method to constrain the coherence of functional classes while the variance in the classwise fitCons scores is maximized, but it could support the use of much larger, richer sets of covariates.

We have focused on HUVEC in this paper, but we also generated fitCons scores for two other cell types (H1 hESC and GM12878). A comparison across cell types (see Supplementary Note) indicated that the genomic positions assigned to each functional class differed substantially across cell types, but equivalently defined clusters had highly similar fitCons scores in the different cell types (Supplementary Fig. S5). When cell-type-specific scores were examined, elements “active” in that cell type displayed significantly higher scores than inactive elements. Moreover, particular elements had higher scores in cell types for which they were active than in cell types for which they were inactive (Supplementary Fig. S6).

This property of cell-type-specificity in the scores is useful in some cases, but in others it is desireable to have a single set of scores that integrate information from multiple cell types. We found that a set of “integrated” scores based on a simple, heuristic procedure (see Methods) performed nearly as well as the cell-type-specific scores in the target cell types, but much better on elements from mismatched or pooled cell types (Supplementary Fig. S7). With more powerful clustering techniques, it may be possible to improve these methods by considering all cell types simultaneously, clustering sites by multi-cell-type functional genomic fingerprints, and then producing a single set of scores reflecting these joint patterns. In this way, nucleotide sites that not only exhibit similar functional genomic fingerprints, but that are active in similar cell types, would be grouped together. Improvements in clustering, together with steady improvements in the resolution and quality of the available functional genomic data, should result in improved power for individual functional elements and refined estimates of the share under selection.

## Acknowledgements

We thank Leonardo Arbiza for helpful discussions and assistance with early analyses, and Greg Cooper for constructive criticism of our validation experiments and comparisons with CADD. This research was supported by NIH grant GM102192, a David and Lucile Packard Fellowship for Science and Engineering (to AS), and a postdoctoral fellowship from the Cornell Center for Comparative and Population Genomics (to IG).

## Online Methods

### Functional Genomic Data

RNA-seq and DNase-seq data for HUVEC, H1 hESC, and GM12878 were downloaded from the University of California, Santa Cruz (UCSC) Genome Browser (http://genome.ucsc.edu). Chromatin states for the same three cell types were downloaded from the European Bioinformatics Institute ftp site (see Supplementary Table S3). For DNase-seq, we considered two replicate experiments from University of Washington (UW) data for each cell type. However, only one UW replicate was available for H1 hESC so additional DNase-seq data for this cell line was obtained from Duke University. For each replicate experiment, we downloaded broad and narrow peak calls. For RNA-seq, we selected Caltech poly-A+ 75 bp paired-end read data, after examining several data sets and considering trade-offs among data quality, coverage and read depth. A single RNA-seq replicate experiment was used per cell type. For chromatin states, we used the 25-state ChromHMM segmentation generated in December, 2012^33^. In addition, coding sequences (CDS) were identified uniformly across cell types using GENCODE data (see below).

### Clustering Approach

We produced a separate partitioning for each cell type based on the functional genomic data. The broad and narrow DNase-seq peaks were used to partition sites in the genome into the following three mutually exclusive classes: sites that fall in a narrow peak in both replicate experiments (2); sites that fall in a broad peak in at least one of the two replicates and do not fall in a narrow peak in both replicates (1); and sites that fall outside of all called peaks (0). This three-level scheme was designed to allow for both high sensitivity (class 1) and high specificity (class 2). For H1 hESC, only one set of broad peak calls was available to define class 1. For the RNA-seq data, we partitioned sites in the genome into four mutually exclusive classes (0–3) based on numbers of reads aligned at each position. Read depth thresholds were set separately for each cell type through a process that aims to minimize the conditional entropy of concentrations of predicted sites under selection (see **Supplementary Note**). The thresholds chosen, in RPM units, were (1, 7, 36) for HUVEC, (1, 5, 20) for H1 hESC, and (1, 11, 38) for GM12878. Chromatin states were defined directly from the 25 states in ChromHMM, except that a 26th state was defined for sites not assigned to a chromatin class. The Cartesian product of these partitions, together with the partition into coding and noncoding sequences (see below), resulted in 3 *×*4*×*26*×*2 = 624 distinct functional classes.

### Running INSIGHT

INSIGHT was used to compute the fitCons score for each non-empty site cluster (see Table S1). The INSIGHT method infers the fraction of nucleotide sites under selection (*ρ*) for a given collection of sites by comparing patterns of within-species polymorphism and between-species divergence for that collection with the patterns observed in putatively neutral sites nearby. A detailed description of the method, sequence data, and data-quality filters is given in reference [34]. Briefly, we used high-coverage genome sequences for 54 unrelated human individuals from the 69 sequences released by Complete Genomics (http://www.completegenomics.com/public-data/69-Genomes/)^50^, as well as the chimpanzee (panTro2), orangutan (ponAbe2), and rhesus Macaque (rheMac2) reference genomes. Various filters were applied to eliminate repetitive sequences, recent duplications, CpG sites, and regions not showing conserved synteny with out-group genomes. For each fitCons site cluster, INSIGHT was used to compute a maximum likelihood estimate of *ρ*, as well as a standard error approximated using the curvature method. The estimate of *ρ* was used as the fitCons score associated with all sites in that cluster. To ensure that our results were not influenced by estimates with high uncertainty, we filtered out clusters for which the estimated standard error was larger than 40% of the estimated value of *ρ*. To increase computation efficiency, clusters larger than 20 Mb were partitioned into smaller subclasses, and INSIGHT was applied separately to each of these subclasses. An estimate of *ρ* for the entire cluster was obtained by taking the average of subclass estimates of *ρ* weighted according to the number of unfiltered sites in each sub cluster. The standard error of an estimate of *ρ* averaged across several site clusters was computed by taking the square root of the weighted mean of the site cluster standard errors squared.

### Neutral Sites

The collection of sites predicted to be free from the influence of natural selection (“neutral” sites) used both by INSIGHT and in some our power analysis (see below) was derived from a set identified previously^28, 34, 51^. Briefly, this set is obtained by starting with all sites in the genome and eliminating several classes of sites likely to be under direct natural selection, including (1) exons of annotated protein-coding genes and the 1000 bp flanking them on either side; (2) RNA genes from GENCODE v11 and 1000 bp flanks; and (3) conserved non-coding elements (identified by phastCons) and 100 bp flanks. We applied the quality filters described above to these sites as well. INSIGHT matches positions in site clusters with putatively neutral sites using a 10 kb sliding window, taking care to avoid matching sites on opposite sides of a recombination hotspot.

### GENCODE Annotations

Transcript annotations from GENCODE v15^52^ were downloaded from (ftp://ftp.sanger.ac.uk/pub/gencode/release15/gencode.v15.annotation.gtf.gz), and used to define eight site classes: coding sequences (CDS), 5′ untranslated regions (UTRs), 3′ UTRs, promoters, introns, long intergenic noncoding RNAs (lincRNA), short noncoding RNAs (sncRNA), and intergenic (sites not falling within any protein-coding transcription unit). Transcripts annotated with feature_type=“CDS” and gene_type=“protein_coding” were used to define the CDS set for fitCons. For subsequent analysis, we used a slightly more conservative set, obtained by additionally requiring feature_type=“gene”, gene_status=“KNOWN”, transcript_status=“KNOWN”, and the identification of both start and stop codons within the transcript. UTRs were defined from transcripts having feature_type=“UTR” and gene_type=“protein_coding”. Each UTR was designated as 5′ or 3′ based on whether it was immediately upstream of the start codon or immediately downstream of the stop codon, respectively. Introns were defined by positions that fall within a protein-coding transcript but outside of the CDS and UTR regions. Promoters were defined as the 1000 bp immediately upstream of the first (i.e., most upstream) transcription start site for each protein-coding gene. A similarly defined alternative set of 100 bp promoter regions was used in assessing differences between cell types (see Fig. S6). LincRNAs were identified by transcripts with feature_type=“exon” and gene_type=“lincRNA”. Similarly, sncRNAs consisted of transcripts with feature_type=“exon” and gene_type ∈ {“miRNA”, “snRNA”, “snoRNA”}. Positions in the more inclusive CDS set were removed from all noncoding classes. When computing the composition of sites exceeding various fitCons score thresholds by annotation type (Fig. 2A), if multiple annotations applied to a nucleotide position, it was assigned to a single category in the following order of precedence: CDS, TFBS, promoter, sncRNA, lincRNA, 5′ UTR, 3′ UTR, intron, and intergenic.

### *Cis*-Regulatory Elements

Transcription factor binding sites (TFBSs) were drawn from a set for 78 transcription factors, based on chromatin immunoprecipitation and sequencing (ChIP-seq) data from ENCODE28 (downloadable at (http://genome-mirror.bscb.cornell.edu/). This set contains roughly 1.4 million binding sites of mean length 11 bp, each of which is associated with the cell types in which it was detected. For some tests, we considered only the subset of nucleotide positions inside these TFBSs that corresponded to motif positions with strong base preferences, defined as those positions at which the consensus allele appeared in at least 90% of all binding sites (according to the inferred motif model). For enhancers, we used the distal regulatory modules described in reference [37]. We downloaded the file enets4.Distal cell line.txt from http://encodenets.gersteinlab.org/ and extracted from it a total of 19,005 enhancer-transcript associations, covering 5,834 unique autosomal loci with a mean length of 888 bp, along with the cell types associated with each predicted enhancer. Expression quantitative trait loci (eQTL) described in reference [6] were downloaded from www.ebi.ac.uk/arrayexpress/files/E-GEUV-1/analysis results/.

We used the four files {EUR373,YRI89}.{exon,gene}.cis.FDR5.best.rs137.txt.gz to identify 6,760 distinct autosomal positions and the associated transcripts. As with other noncoding classes, we removed all positions overlapping CDS.

### Identifying Active Elements per Cell Type

In several analyses, we considered the subset of elements in each annotation class for which we had evidence of activity in a given cell type. To identify the cell types in which TFBS and enhancers were active, we simply used the cell type designations provided in the corresponding annotation files (see above). For other classes of elements, e.g., eQTLs and promoters, we defined the active elements using a set of GENCODE transcripts and genes that showed significantly elevated levels of RNA transcription in the Caltech RNA-seq data. For this purpose, we downloaded from the UCSC Genome Browser files containing normalized read counts in reads per kilobase per million at the levels of both transcripts and genes for HUVEC, H1 hESC, and GM12878 (http://hgdownload.cse.ucsc.edu/goldenPath/hg19/encodeDCC/wgEncodeCaltechRnaSeq/). A transcript (or a gene) was defined as being active in a cell type if the 95% confidence interval if its normalized read count in that cell type fell within the top one third of normalized read counts for transcripts (or genes). The threshold determining the top one third (1.477 for transcripts and 4.966 for genes) was computed by aggregating information from all three cell types. We then determined the set of active eQTLs in each cell type as the ones associated with an active gene, using the GENCODE gene identifier specified for each eQTL in the data file. Similarly, we defined elements in our collections of promoters, UTRs, CDS, and introns, as active if they were associated with an active transcript. For the comparison between cell types (Fig. S6) we also used collections of eQTLs and promoters found to be inactive in a given cell type. Those were defined in a similar way, by using transcripts and genes falling in the bottom third of the distribution of normalized read counts.

### Comparison with Other Scores

Base-wise scores from the Genomic Evolutionary Rate Profiling (GERP)13 method were downloaded from http://mendel.stanford.edu/SidowLab/downloads/gerp/ (file hg19.GERP scores.tar.gz, generated in August, 2010). Scores from phastCons^12^ and phyloP^15^ for 46 placental mammals were downloaded from the UCSC Genome Browser (http://hgdownload. cse.ucsc.edu/goldenPath/hg19/; subdirectories phastCons46way/placentalMammals/ and phyloP-46way/placentalMammals/). The Combined Annotation Dependent Depletion (CADD) scores^35^ were downloaded from http://cadd.gs.washington.edu/download/ (file whole genome SNVs.tsv.gz, downloaded in September, 2013). This file specifies for each genomic position a separate score for each of the three possible variant bases in that site. We took the maximum of these three scores, which yielded the best performance for CADD in our comparisons. We also considered RegulomeDB^36^ as a method for ranking single nucleotide polymorphisms (SNPs), such as eQTLs, according to evidence from functional genomic data. RegulomeDB classifies each common SNP into one of 13 categories (1a–f, 2a–b, 3–7) ranked from strongest (1a) to weakest (7) evidence for function. We downloaded the thirteen categories from http://regulome.stanford.edu/downloads/ in January, 2013.

### Receiver Operating Characteristic (ROC) Curves

We used ROC curves to describe the ability of each scoring scheme to discriminate between functional and nonfunctional TFBSs, enhancers, and eQTLs. For TFBSs and enhancers, we used annotations to indicate the set of functional elements (see above) and used as a non-functional control our filtered, putatively neutral sites (see above). For eQTLs, our control set consisted of all 9.8 million variants tested in reference [6], excluding indels and non-simple variants, and positions that showed possible associations at a threshold of nominal *p <* 0.05 (7.6 million SNPs remained). In all three cases, we removed any sites in our functional set from the negative control set. For each scoring scheme and annotation type, a point on a ROC plot indicates the fraction of annotated genomic positions with scores higher than a given score (true positive rate) versus the fraction of control genomic positions with scores higher than that score (false positive rate). In computing the fractional coverage for each scoring scheme, we ignore positions that are not scored, so as not to penalize methods such as RegulomeDB that provide partial coverage. Note that the other four scoring schemes had similar overall coverage to one another.

### Integrating fitCons Scores Across Cell Types

The main challenge in generating fitCons scores that integrate functional genomic data across cell types, within the context of our simple partitioning scheme, is avoiding a combinatorial explosion in the number of functional genomic clusters considered. We addressed this problem by fixing the partitioning scheme to the original 624 fingerprints (see above), but altering the rule by which nucleotide sites are assigned to clusters to reflect information from multiple cell types. In particular, we attempted to select, for each nucleotide site, the single cluster—from all clusters to which that site was assigned across cell types—that was likely to be most informative about the site’s function. Toward this end, we computed a cell-type aggregated estimate of *ρ* for each of the 624 classes by running INSIGHT on the collection of *all* sites associated with that class in any of the three cell types. Note that, unlike in the standard fitCons pipeline (see Fig. 1), these collections of sites overlap with one another. We then partitioned the sites into non-overlapping clusters by choosing, for each genomic position, the cluster (out of the three) that had the highest cell-type aggregated *ρ*. Finally, we executed INSIGHT once more on each of these disjoint clusters to obtain cell-type integrated fitCons scores.

### Share Under Selection

Assume a partitioning of the genome into *K* mutually exclusive and exhaustive clusters, *C*_1_,…, *C_K_*, and a corresponding set of fitCons scores, *ρ*(*C*_1_),…, *ρ*(*C_K_*). Note that the expected number of genomic positions under selection in cluster *C_i_* is given by *ρ*(*C_i_*)*|C_i_|*, because *ρ* is an estimate of the fraction of sites under selection. Thus, for an arbitrary collection of sites, *S*, the expected number of sites in *S* that are under selection is given by sel(*S*) = ∑*_i_ρ*(*C_i_*)*|C_i_∩S|*, and the average fitCons score for *S* is given by *ρ*(*S*) = sel(*S*)/*|S|*. To avoid under-estimation of *ρ*(*S*), we do not filter out fitCons scores with high uncertainty in these calculations, as we do for other analyses (see above). In addition, to account for possible over-estimation of *ρ* in very large clusters having low fractions of sites under selection, we ran INSIGHT on the large collection of sites (790 Mb) obtained by intersecting the collection of putative neutral sites used to fit the neutral model in INSIGHT (see above) with the cluster, *C*_null_, consisting of non-coding sites in the ‘Quiescent’ chromatin state having no DNase-seq or RNA-seq signal (0 levels defined above). We then subtracted the estimated value of *ρ*, denoted *ρ*_neut_, from the raw fitCons score to obtain a conservative lower bound, *ρ*(*S*) − *ρ*_neut_, for the fraction of sites under selection in *S*.

### FitConsD and Evolutionary Turnover

Our measures of turnover are based on comparisons between fitCons scores, which estimate the fraction of sites under selection since divergence from chimpanzee, with scores obtained using a parallel method, fitConsD, which measures natural selection in longer evolutionary timescales, namely, since the divergence of human, chimpanzee, orangutan, and rhesus macaque. To make this comparison as direct as possible, fitConsD scores were computed using the same pipeline we developed for fitCons (see Fig. 1), except that in step C we replaced the INSIGHT model with an evolutionary model that considers sequence divergence between the four primate genomes.

Briefly, we obtained these divergence-based estimates as follows. We downloaded the multiple genome alignment for 46 placental mammals from the UCSC Genome Browser (http://genome.ucsc.edu), and extracted from it the subalignment for the four primates. In each of the three non-human genomes, we filtered out nonsyntenic regions and positions with genotype quality below 20. Additionally, we masked out sites filtered in the INSIGHT analysis to eliminate repetitive sequences, recent duplications, and CpG sites (see above). We assumed a fixed branch-weighted phylogeny *T* for the four-species tree, which was obtained by fitting a phylogenetic substitution model to fourfold degenerate sites in coding sequences, and we used the partitioning of the genome into 624 clusters {*C_i_*} defined using functional data from the HUVEC cell line (see above).

With these preparations, we estimated a divergence-based fraction of sites under selection, *ρ*_div_(*C_i_*) for each cluster *C_i_*, as follows. First, we created a pseudo-alignment consisting of the columns from the original four-species alignment that correspond to positions in *C_i_*. We then used the *phyloFit* procedure from RPHAST^53^ to estimate a maximum-likelihood scaling factor *s_i_* for the tree *T* for this pseudoalignment. This scaling factor *s_i_* is an estimate of the relative evolutionary rate in cluster *C_i_*, compared with the pre-estimated neutral model, but it does not yet consider variation in the neutral substitution rate along the genome. Therefore, we additionally computed similar scale factors for 10 kb blocks of neutral sites across the genome, using the same neutral sites and windowing scheme as for the fitCons scores (see above). Specifically, for each 10 kb window *w*, we computed a maximum-likelihood neutral scaling factor 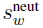 for *T*. We then defined the neutral scale factor 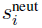 for a cluster *C_i_* as the weighted average of neutral scale factors 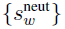 in the associated neutral blocks (i.e., the average is weighted by the size of the intersection and each window *w*). Now the relative rate of substitution in *C_i_* compared to the expectation under neutrality could be computed as, 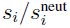. Under the assumption that negative selection dominates^54^, an estimate of the fraction of sites under selection in cluster *C_i_* is therefore given by 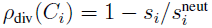.

## Supplementary Tables

**Supplementary Table S1.** FitCons Site Clusters.

| Cell type | Number of clusters |  |  | Coverage<sup>d</sup> | Median size<sup>e</sup> |
| --- | --- | --- | --- | --- | --- |
|  | Non-empty <sup>a</sup> | Viable <sup>b</sup> | Confident <sup>c</sup> |  |  |
| HUVEC | 560 | 502 | 295 | 99.5% | 120 kb |
| H1 hESC | 557 | 498 | 298 | 99.1% | 121 kb |
| GM12878 | 556 | 496 | 287 | 99.4% | 144 kb |
| Integrated | 494 | 466 | 283 | 98.9% | 198 kb |

<sup>a</sup>Number of non-empty clusters in partitioning.
<sup>b</sup>Number of non-empty clusters after applying INSIGHT filters.
<sup>c</sup>Number of clusters for which the ratio between the estimated standard error for $\rho$ and the estimated value of $\rho$ is at most 0.4.
<sup>d</sup>Percentage of bases in confident clusters out of 2,881,033,286 bp in human autosomes.
<sup>e</sup>Median size (in kilobases) of confident clusters.

**Supplementary Table S2.** Share Under Selection for Various Annotation Classes.

| Class | Size (Mbp) | % genome | % sites<br>under sel.<sup>a</sup> | % sel.<br>sites<sup>b</sup> | Adj. % sel.<br>sites<sup>c</sup> |
| --- | --- | --- | --- | --- | --- |
| genome | 2,881.0 | 100.0% | 7.5%±0.1% | 100.0% | 100.0% |
| coding | 30.7 | 1.1% | 63.3%±0.4% | 9.0% | 15.3% |
| 3' UTR | 24.7 | 0.9% | 19.5%±0.3% | 2.2% | 3.3% |
| 5' UTR | 4.1 | 0.1% | 13.3%±0.5% | 0.3% | 0.3% |
| promoter | 21.5 | 0.7% | 9.0%±0.3% | 0.9% | 1.0% |
| ncRNA<sup>d</sup> | 8.0 | 0.3% | 8.1%±0.1% | 0.3% | 0.3% |
| intron | 1,008.8 | 35.4% | 7.5%±0.1% | 35.2% | 35.1% |
| intergenic | 1,768.2 | 61.5% | 6.3%±0.1% | 51.7% | 44.1% |

<sup>a</sup>Fraction of sites under selection in a class computed using HUVEC fitCons scores (see Methods).
<sup>b</sup>Fraction of total number of sites under selection in the genome that is estimated to fall in class.
<sup>c</sup>Fraction of sites under selection estimated to fall in class, corrected by subtracting $\rho_{\text{neut}} = 3.3\%$ from the raw estimate (see Methods).
<sup>d</sup>Union of lincRNA set and sncRNA set.

**Supplementary Table S3.**
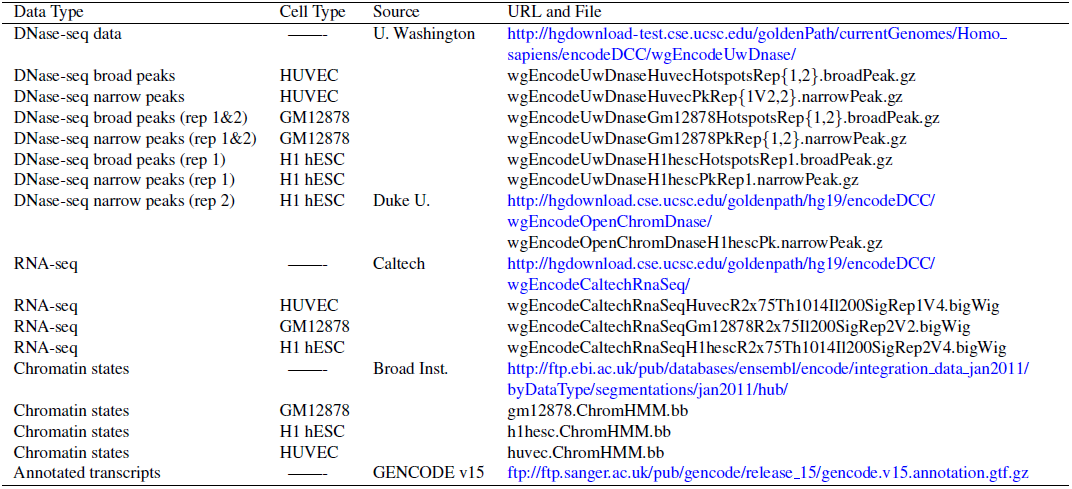
Sources of Functional Genomic Data.

## Supplementary Methods

### Partitioning Genome Based on RNA-seq Data

The DNase-seq and histone modification data provided a natural partitioning of the genome into a small number of classes: using broad and narrow peak calls for DNase-seq, and the 25 ChromHMM states for the histone ChIP-seq data. Using the RNA-seq data to partition the genome, however, required developing a framework that would allow us to determine how informative a given partition is on the distribution of sites under selection in the genome. We used the measure of *mutual information* from information theory, and applied an exhaustive search to find the most informative partition. This exhaustive search was carried out by dividing the range of continuous values (normalized read depth in the case of RNA-seq) into a discrete set of intervals, and assessing the fraction of sites under selection in each interval using INSIGHT. This approach provides a relatively general framework for using fitCons scores computed by INSIGHT to refine a given clustering scheme (see backward arrow from C to B in Fig. 1), and we imagine it will be useful in parsing other complex data sets. The three sections below describe the information theoretic concepts used in our approach, the implementation details of the exhaustive search, and the results for the RNA-seq data for the three cell types.

### Mutual Information and Conditional Entropy

Let *X* be a binary variable indicating whether or not a genomic position is under selection; that is, if a mutation at that site will influence fitness then *X* = 1 and otherwise *X* = 0. In addition, let *Y_C_* indicate the cluster to which the same position is assigned in a given partitioning *C* (*Y_C_* ∈ *C* = {*C*_1_*, …C_K_*). Assuming that sites are selected uniformly at random from the genome and that *ρ*(*C_i_*) denotes the fraction of sites under selection in cluster *C_i_*, the joint probability distribution of *X* and *Y_C_* is given by:
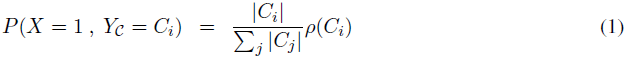
For notational simplicity below, let 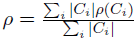, the fraction of sites under selection in the genome.

The mutual information of *X* and *Y_C_* is given by the following expression (Cover and Thomas, 1991):
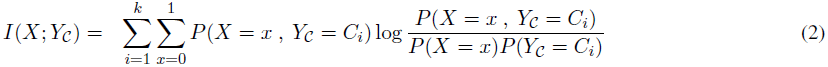

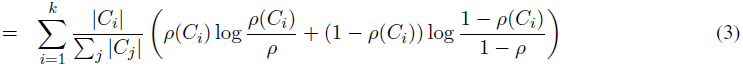

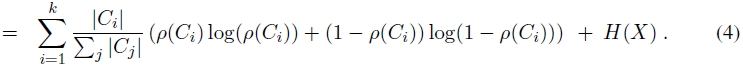
Note that *H*(*X*) = *ρ* log(*ρ*) + (1 − *ρ*) log(1 − *ρ*), the entropy of *X*, does not depend on the partitioning *C*, and the remaining terms on the right hand side of equation (4) are equal to −*H*(*X|Y_C_*), where *H*(*X|Y_C_*) denotes the conditional entropy of *X* given *Y_C_*. Thus maximizing the mutual information of *I*(*X*; *Y_C_*) is the same as minimizing the conditional entropy *H*(*X|Y_C_*).

### Implementation

Our method for partitioning the genome into *K* read-depth bins (for a given *K*) is based on an exhaustive search of all *K*-partitions, *C*, to find the one that results in the largest mutual information *I*(*X*; *Y_C_*). To make the exhaustive search tractable, we apply it to discretized partition boundaries using the procedure outlined below:

1. Divide the continuous range of values (normalized RNA-seq read depth in our case) into *N* discrete intervals, *I_i_,…, I_N_*, such that intervals are of comparable size and large enough to produce confident estimates of *ρ* using INSIGHT.
2. Run INSIGHT on the collection of sites corresponding to each interval *I_i_* to obtain an estimate, *ρ*(*I_i_*), of the fraction of sites under selection in *I_i_*.
3. For each of the 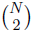 ordered pairs 1 ≤ *i* < *j* ≤ *N*, denote by *I_i,j_* the union of all *I_k_* such that *k ∈* [*i, j*], and estimate *ρ*(*I_i,j_*) using the weighted average 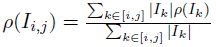
4. For each of the 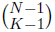 discretized *K*-partitions, *C* = {*C*_1_ *, C*_2_, …, *C_K_*}, defined by *K* + 1 interval boundaries, 0 = *i*_1_ *≤ i*_2_ < *i*_3_ < *… < i_K_ < i_K_*_+1_ = *N*, retrieve for each cluster *C_k_* = *I*_*i*_*k*_ +1,*i*_*k*+1__ an estimate of *ρ*(*C_k_*) from the estimates pre-computed above, and use it to compute the mutual information *I*(*X*; *Y_C_*) using the expression in (4).
5. Choose the *K*-partition with the highest mutual information.

Applying this procedure with increasing values of *K* should result in an increase in the resulting mutual information, but a decrease in the size of clusters.

### Application to RNA-seq data

We applied the procedure described above separately to the RNA-seq data of each of the three cell types. For each cell type, we divided the range of normalized read depth (reads per million; RPM) into *N* = 53 intervals by taking increments of 1 RPM between 0 and 20, increments of 2 between 20 and 40, increments of 5 between 40 and 100, increments of 10 between 100 and 200, and allocating a single interval for RMP *>* 200. We computed *ρ*(*I_i_*) for each of the 53 intervals (step 2 in the procedure described above), and used it to compute estimates of *ρ* for each of the possible 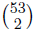 discrete bins (step 3). Then we executed the exhaustive search (steps 4–5) for *K* = 2, 3, 4, 5 (see table below). While the mutual information *I*(*X*; *Y_C_*) kept increasing as we increased *K*, partitioning into more than 4 bins resulted in small bins (less than 30 Mb) for intermediate read depths, which we did not expect to be very informative. We thus chose *K* = 4 for our final partitioning. Note that this partitioning results in one boundary at RPM = 1, another boundary near the deflection point for *ρ*(*I_i_*), and a third boundary around the middle of the dynamic range of *ρ* estimates (see figure below).

Partitioning Based on RNA-seq Read Depth.

Partitioning Based on RNA-seq Read Depth
| cell type | $C^a ; I(X; Y_C)^b$ (K=2) | $C ; I(X; Y_C)$ (K=3) | $C ; I(X; Y_C)$ (K=4)<sup>c</sup> | $C ; I(X; Y_C)$ (K=5) |
| --- | --- | --- | --- | --- |
| HUVEC | {7} ; 0.0117 | {1, 14} ; 0.0133 | {1, 7, 36} ; 0.0138 | {1, 6, 14, 50} ; 0.0140 |
| H1 hESC | {7} ; 0.0116 | {2, 15} ; 0.0132 | {1, 5, 20} ; 0.0139 | {1, 3, 7, 20} ; 0.0141 |
| GM12878 | {20} ; 0.0093 | {1, 28} ; 0.0120 | {1, 11, 38} ; 0.0124 | {1, 10, 28, 75} ; 0.0126 |

<sup>a</sup>Partition $C$ is indicated by $K - 1$ thresholds of RPM from our pre-determined set of 52 boundary points.
<sup>b</sup> $I(X; Y_C)$ is the mutual information of the partitioning $C$ and the distribution of sites under selection, as defined in Equation 4.
<sup>c</sup> $K = 4$ was used for the partitioning in fitCons.

### Controlling for Model Misspecification in INSIGHT

The INSIGHT model makes several simplifying assumptions that could potentially influence its estimates of *ρ*. While these assumptions should not generally bias estimates in any particular direction, the fact that *ρ* is restricted to be positive might lead to a slight bias when estimating *ρ* for site clusters that have a near zero fraction of sites under selection. This slight bias might have a nonnegligible influence on our estimate of the fraction of nucleotides under selection (7.5%), because this estimate is obtained by taking a weighted average of estimates of *ρ* across all clusters, and the terms dominating this average belong to large clusters with very low fractions of sites under selection. To estimate the potential effect of this bias, we ran INSIGHT on the collection of sites that belong to our putatively neutral set and have a null functional fingerprint, i.e., DNase-seq and RNA-seq classes 0, ‘quiescent’ chromatin state, and non-CDS. Our expectation is that INSIGHT should infer *ρ* = 0 for this collection of sites, because it is depleted in functional sites, and more importantly, the putatively neutral sites are used by INSIGHT to define the neutral model. Estimating *ρ* for this very large collection of sites (790 Mb) was done by dividing it into sub-clusters smaller than 20 Mb, running INSIGHT on each sub-cluster and taking the weighted average of the resulting estimates (same approach was used in our main pipeline for large site clusters). The resulting estimate of *ρ*_neut_ = 0.033 was then subtracted from the estimates of each site clusters to obtain a conservative lower bound for *ρ* for that cluster.

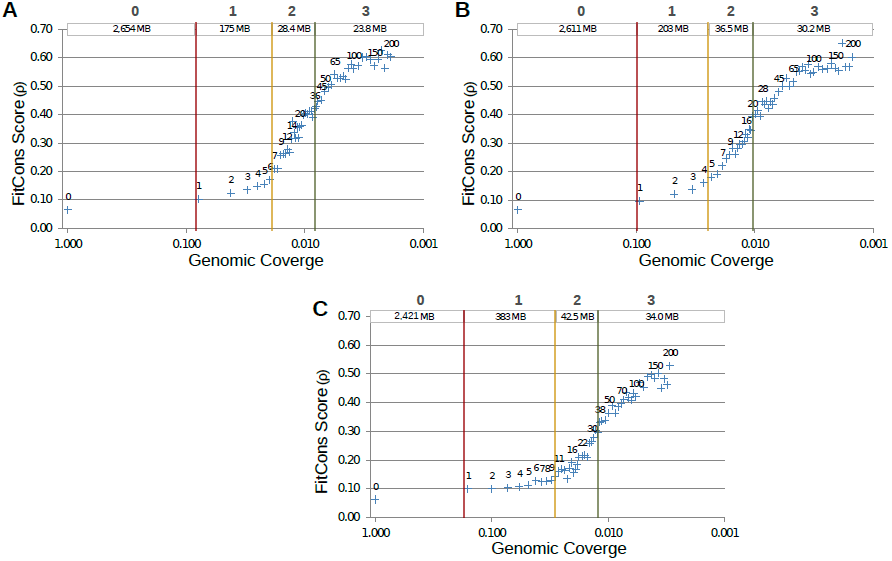
Partitioning the genome into *K* = 4 bins according to normalized read depth in RNA-seq experiments for HUVEC (panel A), H1 hESC (panel B), and GM12878 (panel C). Each point in the scatter plot represents one of *N* = 53 (atomic) intervals *I_i_*, plotting *ρ*(*I_i_*) as a function of the fraction of the genome covered by the union of intervals *I_i_, I_i_*_+1_,…, *I_N_*. The label next to each point corresponds to the lower boundary (in RPM) of that interval. Note that *ρ*(*I_i_*) typically increases with *i*, indicating a higher concentration of sites under selection in highly transcribed sequences. The boundaries between the four resulting classes (0-3) are indicated by vertical lines, with labels (top) representing the class desigbation and the number of position in each class.

### Differences Between Cell Types

Our main analysis focuses on HUVEC, but we also generated fitCons scores for H1 hESC and GM12878. To compare the scores for different cell types, we began by examining the 624 functional genomic classes across the three cell types, in terms of both the genomic positions assigned to each class, and the fitCons scores estimated for those positions. (Note that our partitioning scheme ensures that the the same 624 class definitions are used for each cell type.) Approximately 30% of genomic positions had a null functional fingerprint in all three cell types. In the remainder, we found that genomic positions assigned to each class differed substantially across cell types, with fewer than 4.5% of positions being assigned to the same functional class across all three cell types, and more than a third being assigned to different functional classes in all three cell types. Despite their association with different genomic positions, however, equivalently defined clusters exhibited highly similar fitCons scores across cell types (Pearson correlation *≥* 0.93 for all pairs; Supplementary Fig. S5). Thus, while the patterns of activity differ substantially across cell types, the evolutionary signatures associated with genomic positions that display particular patterns of activity are remarkably consistent across cell types.

**Supplementary Figure S1.**
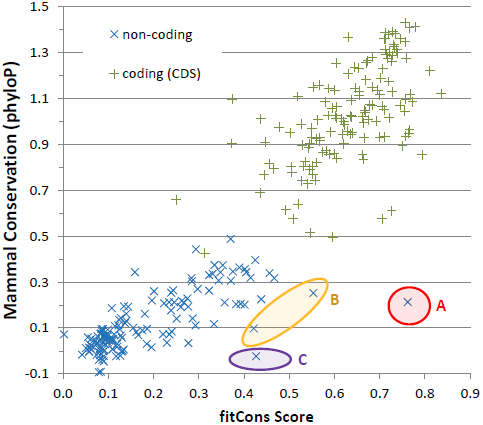
Comparison of fitCons scores and phyloP conservation scores. Each of the 624 clusters is represented by single point, with its *x* coordinate given by the mean placental mammalian phyloP score for the associated genomic positions (Karolchik et al., 2014; Pollard et al., 2010) and its *y* coordinate given by the fitCons score calculated as shown in Fig. 1. The clusters naturally fall in two groups, corresponding to coding sequences (CDS) with lower scores (green crosses) and noncoding sequences with higher scores (blue Xs). Three groups of outliers are shown, representing non-coding clusters with elevated fitCons scores relative to their phyloP scores. Cluster (A) consists of 1200 genomic positions in narrow DNase-seq peaks with no RNA-Seq signal, yet with chromatin modifications indicating transcription activity. These sites are strongly enriched for ChIP-seq-supported TFBSs, and may contain enhancers with weakly expressed eRNAs not detectable from the available RNA-seq data. The two clusters in (B) contain 92.8 kb of sequence defined by high RNA-seq signals, broad DNase-seq peaks, and Pol II binding, and are strongly enriched for 3′ UTR and ncRNA annotations. Cluster (C) contains 52.7 kb of sequence with no DNase-seq but some RNA-seq signal, along with insulator-associated chromatin modifications. This class is strongly enriched for eQTLs and CTCF binding sites, suggesting transcriptional silencing activity. Thus, all four of these clusters appear to be rich in regulatory sequences that could plausibly have experienced weak natural selection during most of mammalian evolution, but come under stronger selection recently on the human lineage.

**Supplementary Figure S2.**
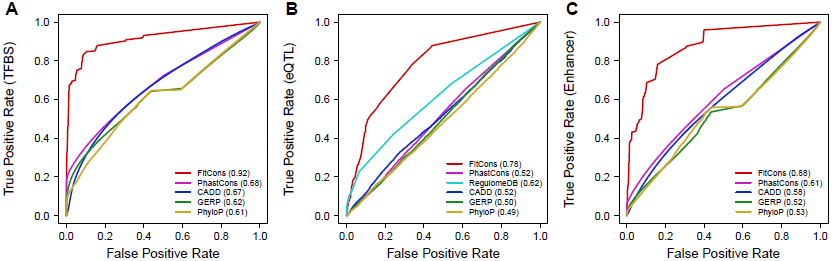
Receiver operating characteristic (ROC) curves for cell-type-specific regulatory elements. Three types of regulatory elements were considered: (A) transcription factor binding sites (TFBSs), (B) expression QTLs (eQTLs), and (C) enhancers identified by chromatin marks. Separate curves are shown for fitCons, phastCons (Siepel et al., 2005), CADD (Kircher et al., 2014), GERP (Cooper et al., 2005), and phyloP (Pollard et al., 2010) scores. In panel (B), a curve is also shown for the RegulomeDB database (Boyle et al., 2012). True positive rates were estimated by the fraction of nucleotides in annotated elements having scores that exceed a given score threshold and false positive rates were estimated by the fraction of nucleotides in a matched set of “null” elements having scores that exceed the same threshold (see Methods for details). Each curve is generated by varying this threshold across the full range of scores for the corresponding method. In this case, only elements “active” in the cell-type for which the fitCons scores were produced (HUVEC) were considered; see Supplementary Fig. S3 for results for a pooled set of elements across cell types. AUC values, shown in parentheses, represent areas under the ROC curve and provide an overall measure of predictive value. The apparent performance of RegulomeDB on eQTLs, particularly at low false positive rates, is somewhat influenced by consideration of eQTL data in its scoring scheme.

**Supplementary Figure S3.**
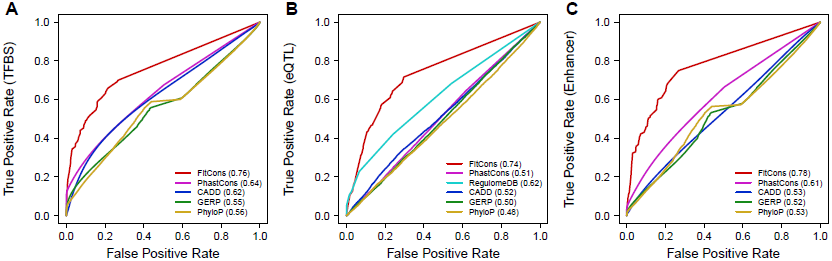
Receiver operating characteristic (ROC) curves for regulatory elements pooled across cell types. Three types of regulatory elements were considered: (A) transcription factor binding sites (TFBSs), (B) expression QTLs (eQTLs), and (C) enhancers identified by chromatin marks. Separate curves are shown for fitCons, phastCons (Siepel et al., 2005), CADD (Kircher et al., 2014), GERP (Cooper et al., 2005), and phyloP (Pollard et al., 2010) scores. In panel (B), a curve is also shown for the RegulomeDB database (Boyle et al., 2012). The fitCons scores used here are computed by aggregating functional information across HUVEC, H1 hESC, and GM12878 cells (see Methods). Note that some regulatory elements might not be active in any of the three cell types. The apparent performance of RegulomeDB on eQTLs, particularly at low false positive rates, is somewhat influenced by consideration of eQTL data in its scoring scheme.

**Supplementary Figure S4.**
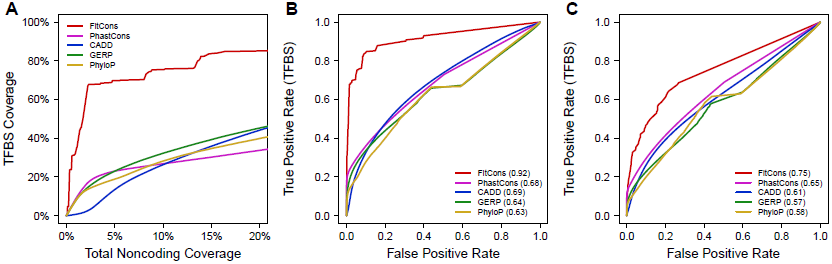
ROC and ROC-like curves for high-information-content positions in transcription factor binding sites. These panels parallel previous figures except that, in this case, only positions in ChIP-seq-annotated transcription factor binding sites with strong nucleotide preferences (relative frequency of preferred allele ≥ 90% in motif model) are considered. Shown are (A) coverage as a functional of total noncoding coverage (as in Fig. 5A); (B) a receiver operating characteristic (ROC) curve for elements active in HUVEC (as in Supplementary Fig. S2A); and (C) a ROC curve based on elements active in various cell types and integrated fitCons scores (as in Supplementary Fig. S3A). These curves show little difference compared with the ones based on whole binding sites, despite known correlations between natural selection and information content for at least some TFs (e.g., see Pollard et al. (2010); Arbiza et al. (2013)), apparently because these correlations tend to be fairly weak and TF-specific, and generally occur below the prediction thresholds of interest.

**Supplementary Figure S5.**
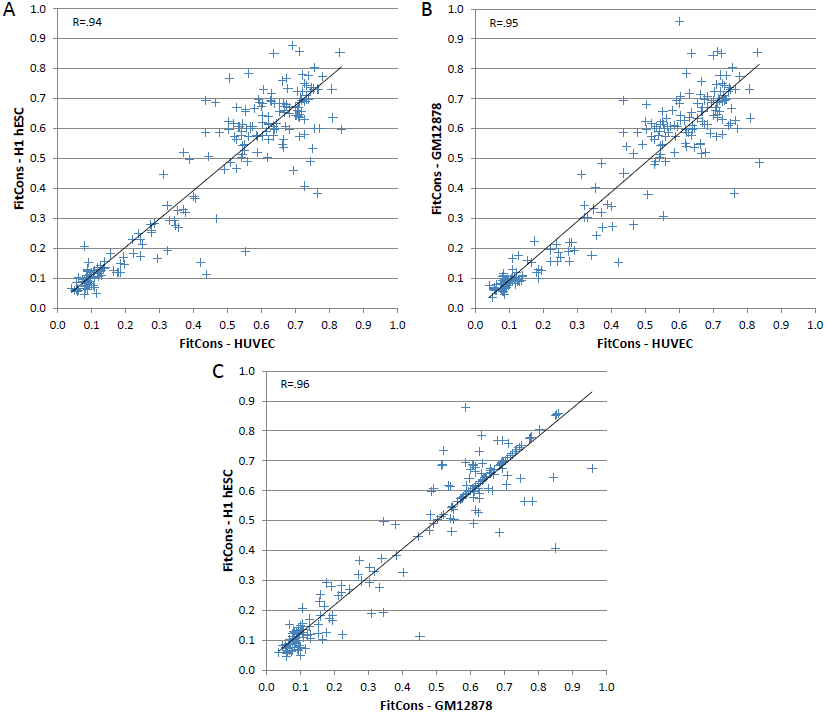
FitCons scores for all functional classes for (A) HUVEC vs. H1 hESC, (B) HUVEC vs. GM12878 and (C) GM12878 vs. H1 hESC cells. While the individual positions assigned to each class vary widely according to cell type, the fitCons scores remain relatively constant, with Pearson correlations *≥* 0.93 and Spearman correlations *≥* 0.87 between pairs of cell types.

To examine the degree to which the scores convey cell-type-specific information, we next considered fitCons scores for elements that are active in one cell type an inactive in another. In particular, we examined subsets of eQTL and proximal promoters (within 100 bp of the annotated transcription start site) that appear to be active in H1 hESC but inactive in HUVEC (H1 HESC+/HUVEC−) or inactive in H1 hESC and active in HUVEC (H1 HESC−/HUVEC+) based on RNA-seq data for the same cell types (see Methods). For each of these groups of elements, we compared mean fitCons scores computed for each of the two cell types (H1 hESC and HUVEC). We found that, based on the scores computed for each cell type, the active elements in that cell type had significantly higher scores than the inactive elements (compare the two gold bars and the two purple bars in each panel in Supplementary Fig. S6). In addition, the same sets of functional elements have significantly higher fitCons scores for the cell type in which they are active than for the one in which they are inactive (compare adjacent gold and purple bars in Supplementary Fig. S6). Similar patterns were observed for comparisons involving GM12878 (results not shown). These findings demonstrate that, while the fitCons scores for all cell types are based on the same polymorphism and divergence data, they nevertheless convey cell-type-specific information through the use of cell-type-specific functional data for clustering.

**Supplementary Figure S6.**
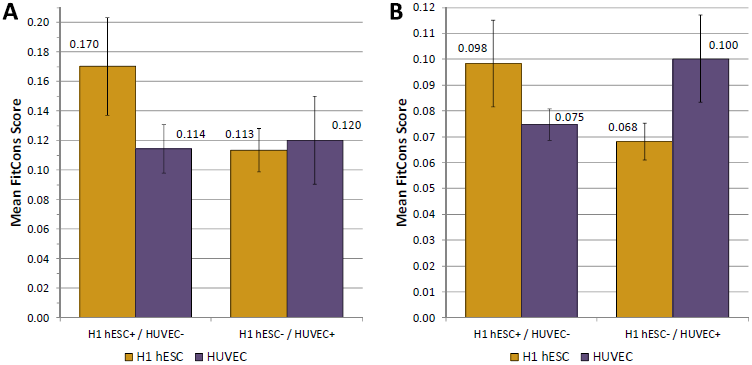
FitCons scores reflect cell-type specific activity. Mean fitCons score for (A) 100 bp promoters and (B) eQTL that are active in one cell type and inactive in another, based on RNA-seq data for the associated gene (see Methods). Error bars represent standard errors of the aggregated fitCons scores (see Methods). FitCons scores computed using functional genomic data from H1 hESC cells (gold bars), elements active in H1 hESC and inactive in HUVEC (H1 hESC+/HUVEC−) are significantly higher than those for elements inactive in H1 hESC and active in HUVEC (H1 hESC−/HUVEC+). The opposite pattern is observed for fitCons scores computed using functional genomic data from HUVEC (purple bars).

**Supplementary Figure S7.**
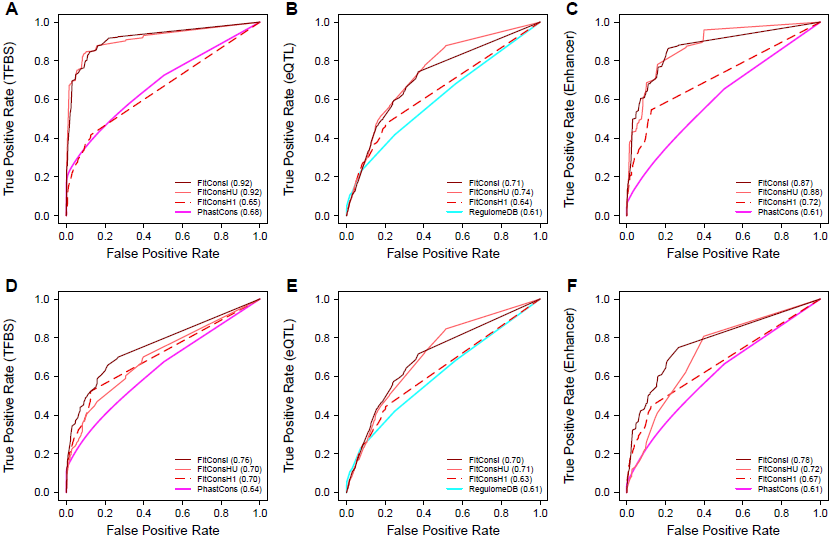
Receiver operating characteristic (ROC) curves comparing integrated fitCons scores with cell-type specific fitCons scores. The top row shows the predictive performance of fitCons scores for elements “active” in the HUVEC cell type: (A) TFBS, (B) eQTL, and (C) Enhancers. Three versions of the fitCons score are shown: cell-type-specific scores based on HUVEC (FitConsHU) and H1-hESC (FitConsH), and scores based on integrated data from all three cell types (FitConsI). Notice that the FitConsI scores perform as well as those based on the “active” cell type (FitConsHU), whereas those based on a different cell type (FitConsH1) perform substantially worse. The bottom row shows the same fitCons scores applied to elements aggregated from a broad range of cell types: (D) TFBS, (E) eQTL, and (F) Enhancers. In this case, FitConsI outperforms both sets of cell-type-specific scores. Thus, the integrated scores (FitConsI) appear to improve performance in a cell-type-general setting without much cost in the cell-type-specific setting.

### Evolutionary vs. Biochemical Measures of “Function”

Following the publications by the ENCODE Consortium in 2012, there has been a great deal of discussion in the scientific literature, the scientific press, and social media about the discordance between evolution-based estimates of the SUS and estimates of the “functional” content of the genome based on high-throughput measures of biochemical activity, which have been reported to be as high as 80% (Dunham et al., 2012; Kellis et al., 2014). For various reasons, the ENCODE-based claims do appear to require a rather generous definition of “function” (Graur et al., 2013; Niu and Jiang, 2013; Doolittle, 2013; Eddy, 2013). Nevertheless, it is worth emphasizing that the question of the functional content of the genome is inevitably dependent on how function is defined.

Consider two possible definitions of “functional” DNA sequences: sequences that produce a phenotype either (1) when mutated (by point mutations), or (2) when deleted. Under the first definition, genomic positions such as fourfold degenerate sites in coding regions or degenerate positions in TFBSs will generally not be functional, whereas under the second definition they will be functional, because their presence is required to maintain the functional coherence of a larger element (they are both examples of “spacer” elements). Other examples of functional sequences whose function does not depend on the precise identity of each nucleotide at each position include sequences separating binding sites for interacting TFs, sequences in short introns, and sequences that maintain the spacing properties of *cis*-regulatory elements relative to target genes.

Importantly, most estimates of the SUS, including ours, have made use of definition (1), whereas measures of biochemical activity are more consistent with definition (2) in some respects (although not all spacer elements will be biochemically active). In our view, it is unlikely that this distinction can account for the difference between estimated genomic fractions of *∼*80% and *∼*5%. Nevertheless, it is worth bearing in mind that our estimate of the SUS and those from comparative genomics are based on a fairly restrictive definition of function. Indeed, our methods indicate that the SUS in annotated coding regions is only about 60%, a fraction that would undoubtedly rise under definition (2).

## References

1. Mardis, E. R. A decade’s perspective on DNA sequencing technology. Nature 470, 198–203 (2011).

2. Wold, B. & Myers, R. M. Sequence census methods for functional genomics. Nat. Methods 5, 19–21 (2008).

3. Dunham, I. et al. An integrated encyclopedia of DNA elements in the human genome. Nature 489, 57–74 (2012).

4. Shen, Y. et al. A map of the cis-regulatory sequences in the mouse genome. Nature 488, 116–120 (2012).

5. Neph, S. et al. An expansive human regulatory lexicon encoded in transcription factor foot-prints. Nature 489, 83–90 (2012).

6. Lappalainen, T. et al. Transcriptome and genome sequencing uncovers functional variation in humans. Nature 501, 506–511 (2013).

7. Cooper, G. M. & Shendure, J. Needles in stacks of needles: finding disease-causal variants in a wealth of genomic data. Nat. Rev. Genet. 12, 628–640 (2011).

8. Mayor, C. et al. VISTA: visualizing global DNA sequence alignments of arbitrary length. Bioinformatics 16, 1046–1047 (2000).

9. Margulies, E. H., Blanchette, M., NISC Comparative Sequencing Program, Haussler, D. & Green, E. D. Identification and characterization of multi-species conserved sequences. Genome Res 13, 2507–2518 (2003).

10. Boffelli, D. et al. Phylogenetic shadowing of primate sequences to find functional regions of the human genome. Science 299, 1391–1394 (2003).

11. Ovcharenko, I., Boffelli, D. & Loots, G. G. eShadow: a tool for comparing closely related sequences. Genome Res 14, 1191–1198 (2004).

12. Siepel, A. et al. Evolutionarily conserved elements in vertebrate, insect, worm, and yeast genomes. Genome Res 15, 1034–1050 (2005).

13. Cooper, G. M. et al. Distribution and intensity of constraint in mammalian genomic sequence. Genome Res 15, 901–913 (2005).

14. Asthana, S., Roytberg, M., Stamatoyannopoulos, J. A. & Sunyaev, S. Analysis of sequence conservation at nucleotide resolution. PLoS Comput Biol 3, e254 (2007).

15. Pollard, K. S., Hubisz, M. J., Rosenbloom, K. R. & Siepel, A. Detection of nonneutral substitution rates on mammalian phylogenies. Genome Res. 20, 110–121 (2010).

16. Graur, D. et al. On the immortality of television sets: “function” in the human genome according to the evolution-free gospel of ENCODE. Genome Biol Evol 5, 578–590 (2013).

17. Niu, D. K. & Jiang, L. Can ENCODE tell us how much junk DNA we carry in our genome? Biochem. Biophys. Res. Commun. 430, 1340–1343 (2013).

18. Doolittle, W. F. Is junk DNA bunk? A critique of ENCODE. Proc. Natl. Acad. Sci. U.S.A. 110, 5294–5300 (2013).

19. Eddy, S. R. The ENCODE project: missteps overshadowing a success. Curr. Biol. 23, R259–261 (2013).

20. McDonald, J. H. & Kreitman, M. Adaptive protein evolution at the Adh locus in Drosophila. Nature 351, 652–654 (1991).

21. Fay, J. C., Wyckoff, G. J. & Wu, C. I. Positive and negative selection on the human genome. Genetics 158, 1227–1234 (2001).

22. Andolfatto, P. Adaptive evolution of non-coding DNA in Drosophila. Nature 437, 1149–1152 (2005).

23. Eyre-Walker, A., Woolfit, M. & Phelps, T. The distribution of fitness effects of new deleterious amino acid mutations in humans. Genetics 173, 891–900 (2006).

24. Boyko, A. R. et al. Assessing the evolutionary impact of amino acid mutations in the human genome. PLoS Genet. 4, e1000083 (2008).

25. Wilson, D. J., Hernandez, R. D., Andolfatto, P. & Przeworski, M. A population genetics-phylogenetics approach to inferring natural selection in coding sequences. PLoS Genet. 7, e1002395 (2011).

26. Ward, L. D. & Kellis, M. Evidence of abundant purifying selection in humans for recently acquired regulatory functions. Science 337, 1675–1678 (2012).

27. Khurana, E. et al. Integrative annotation of variants from 1092 humans: application to cancer genomics. Science 342, 1235587 (2013).

28. Arbiza, L. et al. Genome-wide inference of natural selection on human transcription factor binding sites. Nat. Genet. 45, 723–729 (2013).

29. Narlikar, L. et al. Genome-wide discovery of human heart enhancers. Genome Res. 20, 381–392 (2010).

30. Ritchie, G. R., Dunham, I., Zeggini, E. & Flicek, P. Functional annotation of noncoding sequence variants. Nat. Methods 11, 294–296 (2014).

31. Ernst, J. & Kellis, M. Discovery and characterization of chromatin states for systematic annotation of the human genome. Nat. Biotechnol. 28, 817–825 (2010).

32. Hoffman, M. M. et al. Unsupervised pattern discovery in human chromatin structure through genomic segmentation. Nat. Methods 9, 473–476 (2012).

33. Hoffman, M. M. et al. Integrative annotation of chromatin elements from ENCODE data. Nucleic Acids Res. 41, 827–841 (2013).

34. Gronau, I., Arbiza, L., Mohammed, J. & Siepel, A. Inference of natural selection from interspersed genomic elements based on polymorphism and divergence. Mol. Biol. Evol. 30, 1159–1171 (2013).

35. Kircher, M. et al. A general framework for estimating the relative pathogenicity of human genetic variants. Nat. Genet. 46, 310–315 (2014).

36. Boyle, A. P. et al. Annotation of functional variation in personal genomes using RegulomeDB. Genome Res. 22, 1790–1797 (2012).

37. Gerstein, M. B. et al. Architecture of the human regulatory network derived from ENCODE data. Nature 489, 91–100 (2012).

38. Mouse Genome Sequencing Consortium. Initial sequencing and comparative analysis of the mouse genome. Nature 420, 520–562 (2002).

39. Cooper, G. M. et al. Characterization of evolutionary rates and constraints in three mammalian genomes. Genome Res 14, 539–548 (2004).

40. Lindblad-Toh, K. et al. Genome sequence, comparative analysis and haplotype structure of the domestic dog. Nature 438, 803–819 (2005).

41. Lindblad-Toh, K. et al. A high-resolution map of human evolutionary constraint using 29 mammals. Nature 478, 476–482 (2011).

42. Ponting, C. P., Nellaker, C. & Meader, S. Rapid turnover of functional sequence in human and other genomes. Annu Rev Genomics Hum Genet 12, 275–299 (2011).

43. Chiaromonte, F. et al. The share of human genomic DNA under selection estimated from human-mouse genomic alignments. In Cold Spring Harbor Symp Quant Biol, vol. 68, 245–254 (2003).

44. Meader, S., Ponting, C. P. & Lunter, G. Massive turnover of functional sequence in human and other mammalian genomes. Genome Res. 20, 1335–1343 (2010).

45. Smith, N. G. C., Brandstrom, M. & Ellegren, H. Evidence for turnover of functional noncoding DNA in mammalian genome evolution. Genomics 84, 806–813 (2004).

46. Ponting, C. P. & Hardison, R. C. What fraction of the human genome is functional? Genome Res. 21, 1769–1776 (2011).

47. Lunter, G., Ponting, C. P. & Hein, J. Genome-wide identification of human functional DNA using a neutral indel model. PLoS Comput. Biol. 2, e5 (2006).

48. Kellis, M. et al. Defining functional DNA elements in the human genome. Proc. Natl. Acad. Sci. U.S.A. 111, 6131–6138 (2014).

49. Pheasant, M. & Mattick, J. S. Raising the estimate of functional human sequences. Genome Res. 17, 1245–1253 (2007).

50. Drmanac, R. et al. Human genome sequencing using unchained base reads on self-assembling DNA nanoarrays. Science 327, 78–81 (2010).

51. Gronau, I., Hubisz, M. J., Gulko, B., Danko, C. G. & Siepel, A. Bayesian inference of ancient human demography from individual genome sequences. Nat. Genet. 43, 1031–1034 (2011).

52. Harrow, J. et al. GENCODE: the reference human genome annotation for The ENCODE Project. Genome Res. 22, 1760–1774 (2012).

53. Hubisz, M. J., Pollard, K. S. & Siepel, A. PHAST and RPHAST: Phylogenetic analysis with space/time models. Briefings in Bioinformatics 12, 41–51 (2011).

54. Kondrashov, A. S. & Crow, J. F. A molecular approach to estimating the human deleterious mutation rate. Hum. Mutat. 2, 229–234 (1993).

## References

Arbiza L, Gronau I, Aksoy BA, Hubisz MJ, Gulko B, Keinan A, Siepel A. 2013. Genome-wide inference of natural selection on human transcription factor binding sites. Nat. Genet. 45:723–729.

Boyle AP, Hong EL, Hariharan M, et al. (12 co-authors). 2012. Annotation of functional variation in personal genomes using RegulomeDB. Genome Res. 22:1790–1797.

Cooper GM, Stone EA, Asimenos G, Green ED, Batzoglou S, Sidow A. 2005. Distribution and intensity of constraint in mammalian genomic sequence. Genome Res. 15:901–913.

Cover TM, Thomas JA. 1991. Elements of Information Theory. New York, NY, USA: Wiley-Interscience.

Doolittle WF. 2013. Is junk DNA bunk? A critique of ENCODE. Proc. Natl. Acad. Sci. U.S.A. 110:5294–5300.

Dunham I, Kundaje A, Aldred SF, et al. (1196 co-authors). 2012. An integrated encyclopedia of DNA elements in the human genome. Nature. 489:57–74.

Eddy SR. 2013. The ENCODE project: missteps overshadowing a success. Curr. Biol. 23:R259–261.

Graur D, Zheng Y, Price N, Azevedo RB, Zufall RA, Elhaik E. 2013. On the immortality of television sets: “function” in the human genome according to the evolution-free gospel of ENCODE. Genome Biol Evol. 5:578–590.

Karolchik D, Barber GP, Casper J, et al. (25 co-authors). 2014. The UCSC Genome Browser database: 2014 update. Nucleic Acids Res. 42:D764–770.

Kellis M, Wold B, Snyder MP, et al. (30 co-authors). 2014. Defining functional DNA elements in the human genome. Proc. Natl. Acad. Sci. U.S.A. 111:6131–6138.

Kircher M, Witten DM, Jain P, O’Roak BJ, Cooper GM, Shendure J. 2014. A general framework for estimating the relative pathogenicity of human genetic variants. Nat. Genet. 46:310–315.

Niu DK, Jiang L. 2013. Can ENCODE tell us how much junk DNA we carry in our genome? Biochem. Biophys. Res. Commun. 430:1340–1343.

Pollard KS, Hubisz MJ, Rosenbloom KR, Siepel A. 2010. Detection of nonneutral substitution rates on mammalian phylogenies. Genome Res. 20:110–121.

Siepel A, Bejerano G, Pedersen JS, et al. (16 co-authors). 2005. Evolutionarily conserved elements in vertebrate, insect, worm, and yeast genomes. Genome Res. 15:1034–1050.

